# Manipulating synthetic optogenetic odors reveals the coding logic of olfactory perception

**DOI:** 10.1101/841916

**Authors:** Edmund Chong, Monica Moroni, Christopher Wilson, Shy Shoham, Stefano Panzeri, Dmitry Rinberg

## Abstract

How does neural activity generate perception? The spatial identities and temporal latencies of activated units correlate with external sensory features, but finding the principal activity subspace that is consequential for perception, remains challenging. We trained mice to recognize synthetic odors constructed from parametrically-defined patterns of optogenetic activation, then measured perceptual changes during extensive and controlled perturbations across spatio-temporal dimensions. We modelled recognition as the matching of patterns to learned templates, finding that perceptually-meaningful templates are sequences of spatially-identified units, ordered by latencies relative to each other (with minimal effects of sniff). Within templates, individual units contribute additively, with larger contributions from earlier-activated units. Our synthetic approach reveals the fundamental logic of the olfactory code, and provides a general framework for testing links between sensory activity and perception.

## Introduction

A familiar object evokes a complex pattern of activity in the brain, but it is possible that only a structured subset of this activity, representing critical combinations of sensory attributes, is essential for recognition. A key challenge is to identify the subspace of neural activity that induces the percept. This activity may consist of multiple spatial or temporal features, such as which cells respond and when they respond, relative to stimulus onset or each other. Do individual features contribute differentially to the formation of the percept? For example, the activity of some cells in a pattern may be more important than others. Does the formation of the percept depend on how features are combined? The sequential activation of multiple cells or the latency of their activation relative to brain rhythms are examples of feature combinations that may be perceptually meaningful(*1–3*).

The difficulty in addressing these questions is two-fold. First, multiple features co-vary with perceptual responses, making it difficult to disentangle their independent contributions to perception. Previous studies have mainly focused on correlating neural activity with perception, where the contributions of individual features are entangled. Second, we lack a single framework for quantitatively comparing the perceptual contributions of individual and combined features. For example, we know from causal manipulation studies that single neurons (*4*) or small timing differences (*5–7*) can affect perception, but not their relative importance and how they come together in larger patterns to produce perception.

Here we developed a novel framework for finding the perceptually-meaningful spatiotemporal subspace of neural activity. We optogenetically manipulated individual activity features independently of other features, manipulated combinations of features, and compared the effects of all manipulations under a common metric.

We used mouse olfaction as our model system to study these questions, as odor perception is correlated with complex spatio-temporal activity patterns, and these patterns can be optogenetically manipulated in mice while measuring their perceptual responses. Odor-induced activity recruit glomeruli, discrete neuropils in the olfactory bulb which collect receptor inputs from the nose, grouped by receptor type. Spatio-temporal combinations of glomerular activity correspond to unique odorants (*8, 9*). Both the spatial identity or temporal latencies of glomeruli activated in the pattern may be perceptually meaningful(*6, 7*), but the relative importance of each is unknown. It is also unclear what forms a perceptually-meaningful combination of glomerular activation: ordered sequences aligned relative to each other(*10–12*), sequences aligned to sniff rhythm (*6, 7, 13, 14*), or the earliest activated subset of glomeruli within a sniff rhythm (*15, 16*).

To understand how features of glomerular activity combine to produce perception, we first trained mice to recognize synthetic odors: optogenetically-driven spatiotemporal patterns of glomerular activity. We then performed precise spatial or temporal perturbations on trained patterns and measured how recognition changes. Changes in recognition reflect the perceptual relevance of the modified feature (or groupings of features). We modelled recognition as the matching of glomerular activity to learned templates, and uncovered what forms a perceptually-meaningful pattern template: activation sequences ordered by latencies relative to each other (latencies relative to sniff contribute little), with greater perceptual importance carried by the earlier glomeruli in the sequence. Spatially-identified glomeruli within the sequence contribute additively to perception, with minimal interactions between spatial identities. Template matching with these perceptually-meaningful features can account for animals’ responses, with the degree of mismatch predicting changes in recognition. Hence, by performing precise and parametric manipulation of glomerular activity and quantifying effects under a common metric, we derived a unifying model that explains how odor perception arises from structured glomerular activation.

## Results

We first characterized the basic behavioral and neural responses to optogenetic stimulation. We chronically implanted OMP-ChR2-YFP mice (n=6) (*6*) with cranial windows to expose dorsal olfactory bulb, and performed optogenetic stimulation using a digital micromirror device system (Fig. 1A). Similar to previous reports (*11, 17*) and consistent with known anatomy, mitral/tufted (MT) cells were activated by few localized spots (Fig. 1B). We observed instantaneous MT cell firing rates up to ~100 Hz, with excitatory responses lasting around ~80 ms, comparable to odor-evoked responses (*13*). We verified that spots at the same stimulation parameters were perceptually detectable, but only for ChR2-positive mice (Fig. 1C), and without systematic spatial biases (Fig. 1D). We then used the same basic stimulation parameters for the main experiments.

**Figure 1.**
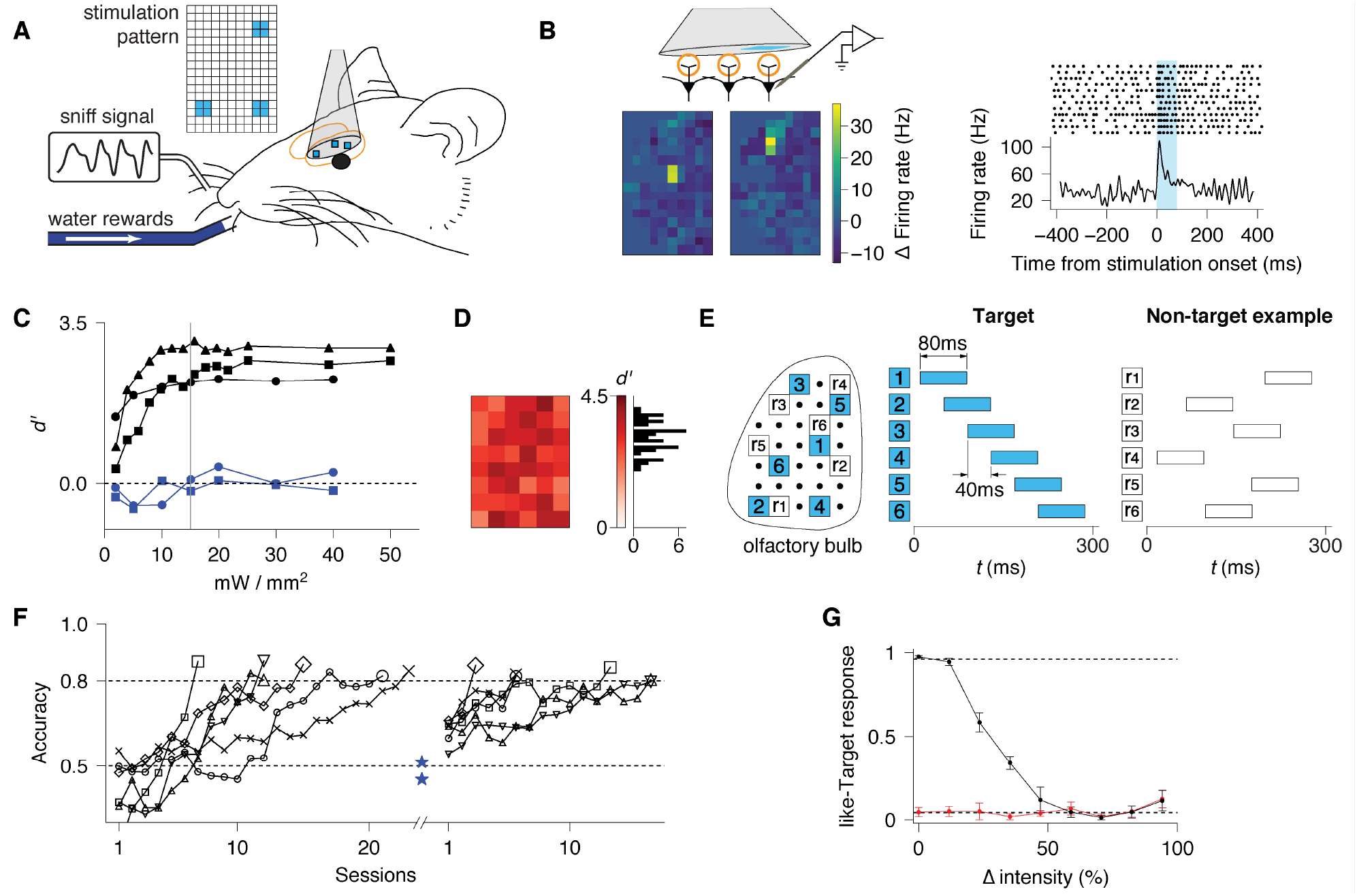
Optogenetic stimulation as synthetic odors. **A**. Schematic of the experimental setup. Dorsal olfactory bulb (OB) was exposed by a chronically-implanted 3 mm window. Spatio-temporal stimulation patterns, created by a digital micromirror device, were projected onto the OB of a head-fixed mouse in front of lick spouts delivering water and a pressure sensor for sniff monitoring. **B.** Mitral/Tufted cell responses to single spot stimulation (120×120 μm^2^, 80 ms duration, 15 mW/mm^2^) across a grid of spots. *Left*: heatmaps of evoked firing (average firing rate across the stimulus duration, 80 ms) for each stimulated spot position, for two typical cells. *Right:* trial-by-trial spiking activity and PSTH corresponding to left cell in (B), for the spot which evoked the largest averaged response. **C.** Behavioral detection *d’* of single spot stimulation at different laser intensities for ChR2-positive animals (black) or ChR2-negative animals (blue). Grey vertical line marks 15 mW/mm^2^. **D.** *d’* for spot detection at 15 mW/mm^2^ stimulation for one ChR2 mouse at different spot positions. Right: Histogram of *d’* across spots. **E.** Schematics for pattern discrimination task. Animals were trained to recognize Target versus Non-target patterns defined on a stimulation grid. Target patterns comprised of six spots, initialized randomly but fixed across subsequent sessions, activated in an ordered sequence defined in time where ‘0’ marks inhalation onset. Non-target patterns were six off-Target spots, randomly chosen from trial to trial, with randomized timing within 300 ms from inhalation (~single sniff). **F.** Learning curves on pattern discrimination task. Mice were initially shaped by discriminating one Target pattern versus one Non-target pattern (left) to criterion performance of 0.8, then trained on the same Target versus multiple Non-target patterns (right). *Note:* performance below chance level (0.5) for some mice at the beginning of training is explained by a strong initial side bias (see Methods). Blue stars indicate performance of control, ChR2-negative mice after >40 sessions of training. **G.** Effect of laser intensity on pattern recognition for mice previously trained to discriminate Target vs Non-target patterns. Probability of like-Target responses (mean and SEM across mice) on probe trials with randomly chosen low intensity Target patterns (black) and Non-target patterns (red). Horizontal dashed lines correspond to baseline responses to Target and Non-target patterns.

We trained mice to discriminate Target versus Non-target synthetic odors on a two-choice task (Fig. 1E). While the stimulation patterns we used are unlikely to correspond to specific known chemical odorants, our patterns follow stimulation durations and activation latencies that fall within known odor-evoked distributions of glomerular activation (*8*). Mice learned to discriminate a fixed Target from any random Non-target pattern over weeks, while ChR2-negative mice did not (Fig. 1F).

We then performed systematic spatial and/or temporal perturbations within Target patterns on a small number (10%) of probe trials, and measured perceptual responses. Perturbations consisted of replacing spots with Non-target spots, or shifting spots in time. For any set of perturbations, we measured the fraction of trials where the mouse makes a lick choice towards the water spout associated with the Target (‘like-Target’ response), as opposed to the Non-target spout. This measurement reflects perceptual distances (*18, 19*): the perceived difference between the perturbed and original Target pattern. The larger the perceptual distance, the lower the fraction of like-Target responses. We performed a basic check that perceptual distances should increase as Target patterns are altered. In pilot mice trained on the task, we systematically decreased the laser intensity below the intensity used during training. As the laser intensity for Target patterns was lowered, animals showed a graded decrease in like-Target responses (Fig. 1G).

We found that spatial perturbations led to graded changes in perceptual distances, dependent on the timing of the spot perturbed. To perform spatial perturbations, we kept Target pattern timing fixed while replacing spots with random Non-target spots (Fig. 2A). One or more spots in Target patterns were replaced in every possible combination of the six spot positions. We found a graded relationship between the number of spots replaced and perceptual distance (Fig. 2B). Spot replacement effects also depended on spot timing: replacing earlier-activated spots had a greater effect than later-activated spots, both within single-spot or multiple-spot replacement trials (Fig. 2C). We refer to this temporal dependence as ‘primacy’ (*15*). We additionally found that the effect of spot replacement depends on spatial distance on the olfactory bulb. The perceptual distance is small between a Target, and a Target with one spot replaced by a proximal spot. This perceptual distance is larger when the new, replacing spot is distant from the old, replaced spot (Fig. S1). This is consistent with studies suggesting that close-by glomeruli may be more likely to be chemically, and hence perceptually, related, at least on a coarse scale (*20, 21*). We cannot completely rule out the alternative possibility that these effects may arise from activating fibres of passage, with strength of activation dependent on distance. However, these effects are likely to be weak given our observed mitral cell responses (Fig. 1B) and also previous reports using similar stimulation parameters (*11*).

**Figure 2.**
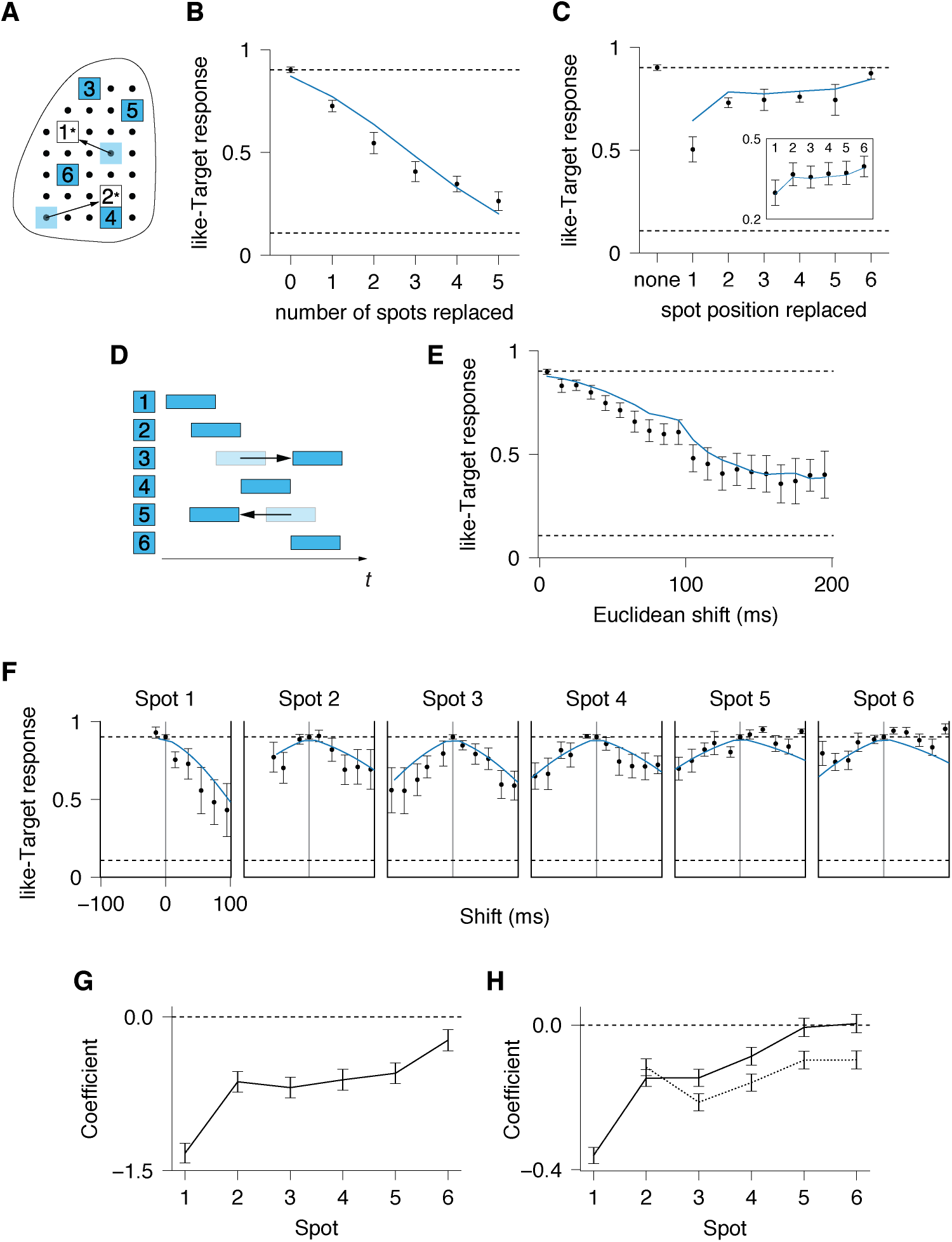
Perceptual effects of spatial and temporal perturbations. **A.** Illustration of spatial perturbations: one or multiple spots in Target patterns were randomly replaced with Non-target spots. **B.** Probability of like-Target responses as a function of number of replaced spots in Target patterns, averaged across animals. Here and later in plots of like-Target responses, horizontal dashed lines correspond to baseline responses to Target and Non-target patterns, and blue lines show the corresponding regression fits. **C.** Probability of like-Target responses as a function of a spot position for single spot replacement. Inset: Probability of like-Target responses as a function of a spot position, marginalized over trials with multiple spots replaced. **D.** Illustration of temporal perturbations, where one or multiple spots in Target patterns were temporally shifted. **E.** Probability of like-Target responses as a function of the amount of temporal shift (‘Euclidean shift’) in the pattern. **F.** Effect of single spot shifts plotted by spot position. **G.** Regression coefficients showing the effect of replacing each spot in all trials, including single or multiple spot replacement. Negative coefficients imply that replacing the spot lowers like-Target responses, coefficients of zero imply no effect on responses. **H.** Same as for (G) but for temporal perturbations. Separate coefficients were fitted for positive shifts (towards the end of the sniff cycle) (solid) and negative shifts (towards the beginning of the sniff cycle) (dashed).

Temporal perturbations also led to graded changes in perceptual distances dependent on the timing of the spot perturbed. To perform temporal perturbations, we shifted one or more spots of the Target pattern in 10 ms increments, in randomly-chosen combinations of spot and shift magnitude (Fig. 2D). We found that increasing the overall amount of temporal shift (defined as 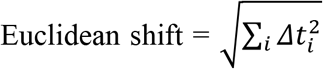 where *Δt*_*i*_ are individual spot shifts) led to an increase in perceptual distance (Fig. 2E). Similar to spatial perturbations, temporal perturbations also exhibit primacy. In trials where only one spot was shifted, shifting earlier-activated spots has stronger effects than shifting later-activated spots (Fig. 2F). Furthermore, we observed an asymmetry in temporal shift effects: shifting spots earlier - towards the beginning of the sniff cycle, has a stronger effect than shifting spots later - towards the end of the sniff cycle. This asymmetry is consistent with primacy: while later-activated spots are less important perceptually, shifting them earlier may interfere with the early, more important spots, hence inducing larger behavioral effects.

To quantify the effect of spatial and temporal perturbations, we performed regression analyses using either binary variables for spot replacement (*x*_*i*_ = 1, if the spot is replaced and *x*_*i*_ = 0, if not), or continuous variables for temporal shifts (*Δt*_*i*_). The regression models quantify primacy in spatial perturbations (Fig. 2G), and primacy and asymmetry for temporal perturbations (Fig. 2H). Primacy is reflected in larger magnitude coefficients for earlier spots; temporal asymmetry is reflected in larger magnitude coefficients for shifting spots earlier as opposed to later. Regression fits are presented in all data plots as blue lines (Figs. 2B,C,E&F), and responses of individual mice are plotted in Fig. S3.

The effects of temporal perturbations suggest that animals encode spot latencies referenced to sniff: as spots were shifted with respect to inhalation onset, animals’ responses changed. An alternative possibility is that spot latencies are encoded with respect to other spots in the pattern. To test this, we considered the synchronous shift of the Target pattern, where all spots are shifted by the same amount with respect to inhalation (Fig. 3A). Synchronous shifts alter sniff-referenced timing while keeping pattern-referenced timing constant. We found that perceptual distances vary, but only weakly, with synchronous shifts (Fig. 3B). Critically, we attempted to predict the effect of any synchronous shift, by summing the effects of single spots shifted by equal amounts referenced to inhalation. The prediction significantly over-estimates the weak perceptual effect of synchronous shifts, suggesting an important role for pattern-referenced timing. Pattern-referenced timing can also be reflected in the order or sequence of spot activation. We found that order-changing temporal perturbations had a larger effect than order-preserving perturbations, controlling for equivalent amounts of sniff-referenced shift from the Target (Fig. 3C). Taken together, the results demonstrate that pattern-referenced timing is a key component of spatio-temporal olfactory activity.

**Figure 3.**
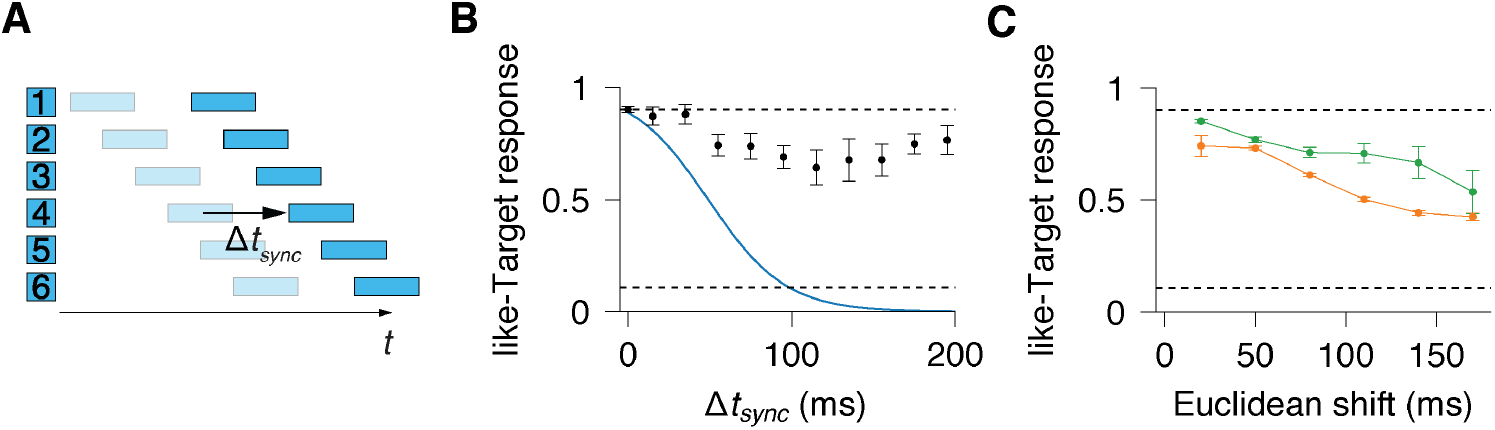
Animals encode pattern-referenced timing. **A.** Illustration of temporal perturbations where Target patterns were shifted synchronously from inhalation by Δ*t*_*sync*_. **B.** like-Target response as a function of Δ*t*_*sync*_ (black), with the predicted response from summing up effects of single spots shifted relative to inhalation (blue). **C.** Comparison of average responses to temporal perturbations which preserve (green) and do not preserve (orange) temporal spot sequence for the same amount of Euclidean shift.

To explain the whole spectrum of behavioral results, we propose a spatiotemporal template matching (STM) model, inspired by template matching models in the broader literature. Template matching is described in psychological theories of pattern recognition as the general process of comparing new inputs to templates stored in memory (*22*). In systems neuroscience, template matching can refer to methods used by experimenters to compare spatio-temporal neural population activity (*23–25*). We propose that olfactory circuits compute similar forms of matching of neural activity for odor recognition.

The STM model consists of input transformation followed by template matching (Fig. 4A). Any olfactory input is transformed to temporal activations in discrete spatial channels (spots, or more generally, glomeruli). Activations decay over time and are represented by exponential waveforms. The decay slope reflects animals’ sensitivity to temporal perturbations, with slower decays corresponding to higher invariance to temporal shifts. The strength of activations is further multiplicatively modulated by primacy, such that earlier-activated channels within the pattern evoke larger waveforms. Each learned target input is transformed and stored as a template which new patterns are compared to.

**Figure 4.**
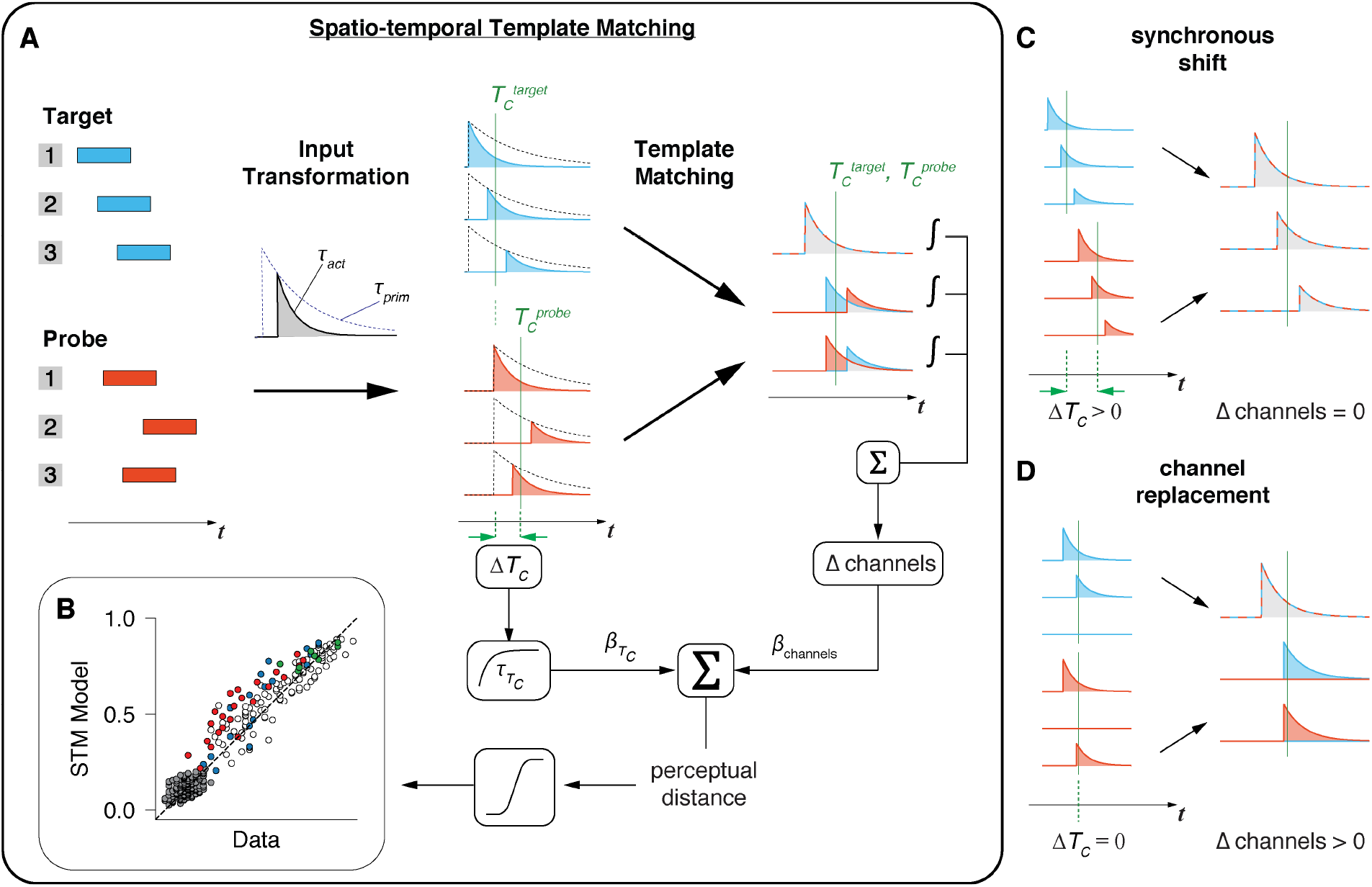
Spatio-temporal Template Matching (STM) maps olfactory inputs to perceptual outputs. **A.** A learned pattern stored in memory (Target, blue) is compared with a new input pattern (Probe, red). Patterns are transformed into waveform representations: pulse activations convolved with exponentials, with decay of activation parameter *τ*_*act*_ and an amplitude, which is a different exponential function (*τ*_*prim*_) of the pulse timing from the onset of the pattern. For each set of waveforms, the center-of-activity 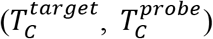 and their absolute difference, 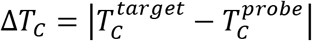, are calculated. Individual waveforms are then aligned to their respective pattern’s center-of-activity, and the difference in non-overlapping area per waveform channel is integrated. This non-overlapping area is summed linearly across all channels (Δ channels). Δ*T*_*C*_ is exponentiated, weighted, then added to the weighted Δ channels value, producing a perceptual distance value. A logistic function on perceptual distances produces a probability of making a like-Target response to the given Probe pattern. **B.** STM model predictions against mouse responses, across all trial types. Within each trial type, trials were sorted by perceptual distance in the model and grouped. Each dot represents a set of 50 trials. The dashed unity line indicates perfect prediction. Different colors indicate different types of perturbation: spatial (blue), synchronous temporal shifts (green), all other temporal shifts (white), Non-targets (grey), spatial and temporal perturbation in same pattern (red). The model has the following fitting parameters: *τ*_*act*_, *τ*_*prim*_, 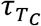, *β*_*channels*_, 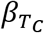 and bias term *β*_*0*_. **C.** Example of template matching for synchronously shifted patterns: only *T*_*C*_ changes, but not waveform differences within channels. **D.** Template matching for patterns where activation in one channel is replaced by another channel: *T*_*C*_ does not change but waveform differences arise.

A template matching procedure then determines the perceptual distance between new patterns and target templates. Template matching considers two effects independently: the difference in the center-of-activity, and differences within individual spatial channels. The center-of-activity captures the average position of each template within a sniff cycle; similar centroid measures have been defined to characterize the mean response latency across multiple neurons (*26*). To perform within-channel comparisons, the model measures the amount of non-overlap between waveforms. Critically, this comparison is done in reference to each template’s center-of-activity. Consider synchronous shifts of Target patterns (Fig. 3B): only the center-of-activity changes, but not waveforms relative to the center-of-activity (Fig. 4C). Hence, within-channel comparisons are primarily sensitive to changes in pattern-referenced timing as opposed to sniff-referenced timing. The choice of using a centroid model for pattern-referenced timing was further corroborated by regression analyses showing that the centroid model best predicts responses to temporal perturbations, compared to other models (Fig. S2). Within-channel comparisons are also sensitive to differences in spatial activations, for example in the spot replacement of patterns (Fig. 4D). The non-overlap per channel is then linearly summed across all channels to produce an overall channel difference value. Channel differences and the difference in center-of-activity are separately weighted, then combined to produce perceptual distances.

The STM model succeeds at predicting animals’ responses to the whole spectrum of pattern perturbations (Fig. 4B) and reproduces the observed trends for individual perturbations (Fig. 5A). The model captures the graded effects of both spatial and temporal perturbations, with primacy in spatial perturbations, and primacy and asymmetry in temporal perturbations. The model also captures the weak effects of synchronous shifts. We further confirmed that changing any component in the model decreases model performance (Fig. 5B).

**Figure 5.**
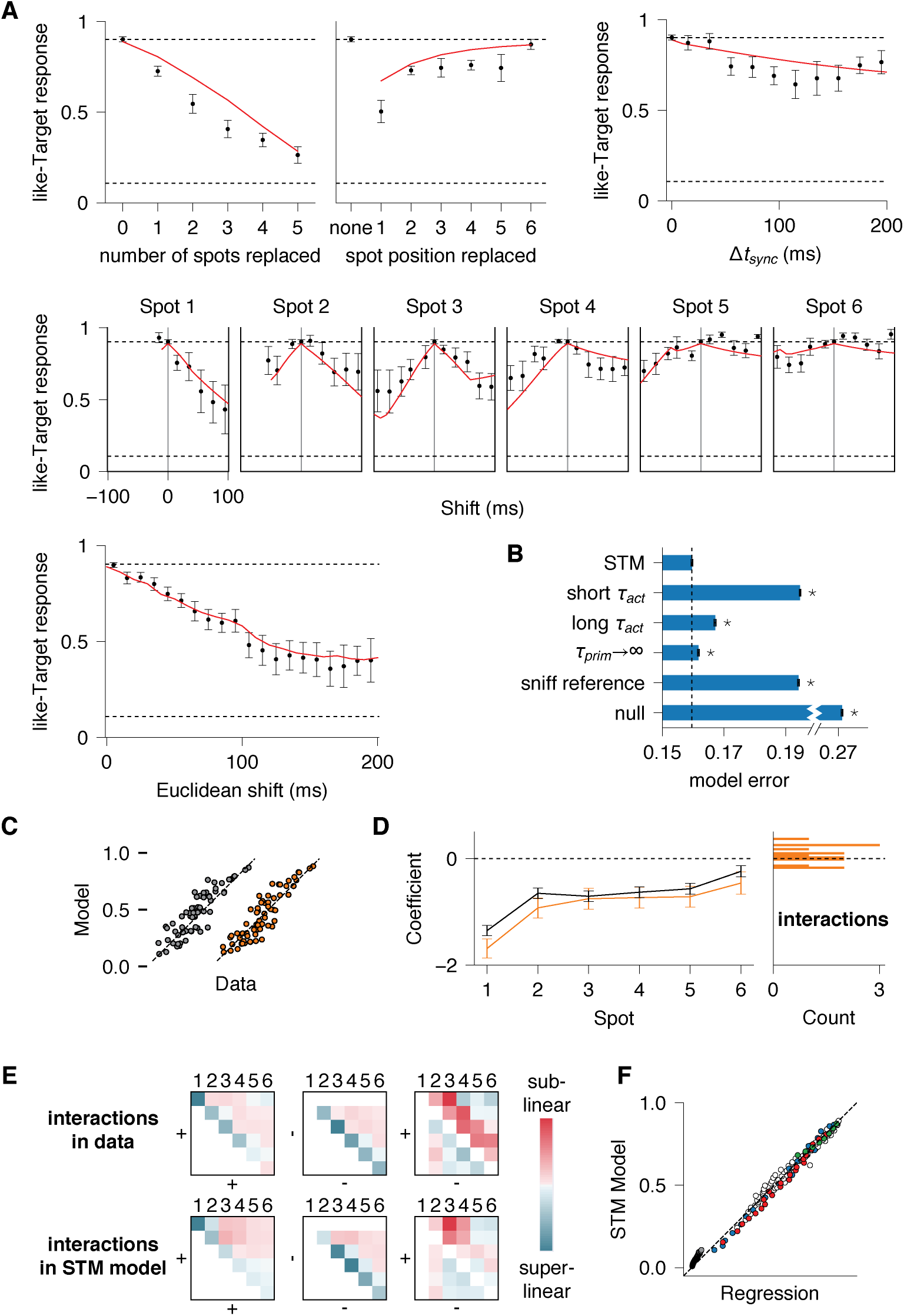
Validation of the STM model. **A.** Fits of the STM model (red) to individual spatial or temporal perturbations. **B.** STM model compared against modified models. Dashed line indicates prediction error of STM Model. ‘short *τ*_*act*_’ and ‘long *τ*_*act*_’: changing *τ*_*act*_ from the fitted value (~60ms) to 10 ms or 200 ms respectively. ‘*τ*_*prim*_→∞’: setting *τ*_*prim*_ to a large value, essentially removing the primacy effect. ‘sniff reference’: Within-channel comparisons referenced to sniff and not center-of-activity. ‘null’: Null model. Asterisks indicate p < .001 significant difference in model prediction error versus STM Model, obtained from ANOVA corrected for multiple comparisons. **C.** Comparison of regression models for spatial perturbations, with non-linear interactions (orange) or without (grey). The unity line (dashed) indicates perfectly predicted responses. Each dot represents a unique combination of spatial replacement by spot position. **D.** Coefficients for models in (**C**), for model with interactions (orange) or without (black), plotted by spot position replaced. Right: histogram of coefficient values for interaction terms. **E.** Comparison of temporal interactions, in data as quantified by regression (top), and in simulated data run through STM model (bottom). We considered pairwise interactions of directional shifts of spots (numbered). Left: shifting two spots later (+,+), Middle: shifting two spots earlier (−,−), Right: shifting one spot later (+) and another spot earlier (−). Colors indicate magnitude and direction of interaction, red / teal indicates sublinear / super-linear interactions, meaning that the effect of paired shift is less / more than predicted from linear sum. **F.** Predictions of STM model plotted against regression models for all data, for the same grouping of trials as in Fig. 4B.

The STM model essentially performs a linear readout of spatial channels, as within-channel differences are summed up linearly without added interactions between channels. If animals respond to patterns according to model predictions, the observed effects of spatial perturbations should also sum linearly. This is because replacing spots while keeping timing fixed, produces only changes within channels (which are summed linearly) (Fig. 4D). We directly tested if a simple linear readout could indeed account for animals’ responses to spatial perturbations. We compared a regression model containing only linear terms, versus a regression model allowing for non-linear effects. The linear model performed better than the non-linear model (Fig. 5C, model error = 0.158 vs 0.16, *p* < 0.001, t-test), and coefficients for non-linear interactions were negligible (Fig. 5D). Hence, the olfactory system performs a linear readout of spatial glomerular activity: glomeruli contribute additively to the overall percept, largely independently of the identity of the other activated glomeruli in the pattern.

On the other hand, the STM model predicts strong temporal interactions between channel activations: shifting any spot changes the latencies of all other spots encoded in reference to the whole pattern. Hence, the effect of shifting any spot depends on how other spots are shifted. For example, the effect of synchronous shifts cannot be predicted from single spot shifts (Fig. 3B). We confirm the existence of temporal interactions with regression: a regression model allowing temporal interactions between spots, is superior to one that does not (model error = 0.1647 vs 0.1725, *p* < .001, t-test). Critically, we found that the STM model run on a large simulated dataset of random temporal perturbations produces the same pattern of temporal interactions as observed in the data (Fig. 5E, pearson correlation = 0.83).

We then quantitatively compared the STM model to the best regression model fitted separately for each individual class of spatial or temporal perturbations. The regression models form a standard for comparison, as they place minimal prior assumptions on the structure of the data. Despite the STM model having fewer parameters than the regressions (6 vs 21), the STM model performs slightly better (prediction error = 0.159 vs 0.160 *p* <.001, t-test). Furthermore, for any given pattern, the STM model and regression models produce responses that are tightly correlated (Fig. 5F); the correlation between models is greater than the correlation between the STM model and the data (*r* = 0.997 vs *r* = 0.97). Hence, the STM model produces similar patterns of responses to regression analyses which could in principle to capture more complex relationships within the data. This implies that the several key computations modelled by STM are sufficient to account for animals’ responses.

We further validated the STM model with unseen classes of spatio-temporal perturbations. In our described experiments and model fitting so far, Target pattern perturbations were either spatial or temporal, but not both. We tested the fitted STM model on additional patterns containing both temporal shifts and spatial replacements within the same pattern. We found that the model could predict animals’ responses to spatio-temporal perturbations despite never encountering these manipulations during model fitting (Fig. 4B, prediction error on spatio-temporal perturbations = 0.23, spatial perturbations alone = 0.23, temporal perturbations alone = 0.21). Another form of spatio-temporal perturbations is inherent in Non-target trials, where all Target spots were replaced and randomly shifted. No Non-target pattern was ever repeated for each animal, guaranteeing that all Non-target stimuli in model fitting are distinct from Non-target stimuli in model testing. The STM model was able to predict responses across different Non-targets (Fig. 4B, error = 0.10).

Hence, spatio-temporal template matching of olfactory bulb activity, linearly summed, constrained by activity-centered timing, and with preferential weighting on early activations, can account for how the olfactory system maps complex activity to perceptual outputs.

## Discussion

We developed a novel framework for finding the perceptually-meaningful subspace of spatio-temporal sensory activity. We optogenetically manipulated features in glomerular activity independently of other features, with manipulations that finely and extensively tile spatio-temporal dimensions, in a model-agnostic manner. By comparing all manipulations under a common metric, we derived a unifying, model-based account of olfactory computations. Hence, while previous studies establish the relevance of single feature changes within patterns (*4, 5, 7*), our approach allows us to understand how features combine in complex patterns to generate perception.

In our study, we chose to measure perceptual distances across manipulations of neural activity. The approach originates from psychology, where the measurement of perceptual distances across manipulations of sensory stimuli (e.g. images, sounds) have revealed underlying models of sensory representation (*18, 19*). By measuring perceptual distances across changes in neural activity, we derived models of neural representation.

The STM model resolves several open issues in olfactory coding. Firstly, the model suggests a linear readout of spatial glomerular patterns. It has been proposed that odors are encoded in unique spatial combinations of glomeruli (*27*), but there are multiple possible combinatorial coding schemes which have not been tested. For example, one extreme possibility is a ‘barcode’ representation, where any slight change to the combination leads to a completely un-related odor, maximizing the system’s representational capacity for odors. Instead, we found a perceptual readout of spatial patterns that is linear. This is consistent with studies which imaged odor-evoked glomerular activity(*28–30*) or directly stimulated individual glomeuruli (*31*). A linear readout implies that two patterns will generate odors that are perceptually similar depending on the degree of glomerular overlap. While a linear readout has lower representational capacity, it may explain the generalization of odor percepts across varying concentrations or backgrounds, where activated glomeruli differ slightly.

Our work causally establishes the role of temporal sequences in odor perception, which has long been hypothesized (*8, 13*) but not directly tested: temporal codes are not theoretically required to support odor perception (*32*). We presently demonstrated that mice trained to recognize a synthetic odor pattern, use temporal sequences in odor recognition even though other (spatial) cues are sufficient to solve the task. Surprisingly, we further found that these sequences were defined with relative latencies within a pattern(*10*), as opposed to latencies with respect to sniff as previously proposed (*6, 8, 13*). Pattern-referenced timing may reflect how inputs compete or are integrated downstream, as competition or integration can depend on temporal proximity between inputs (*11, 33, 34*). We found a weak perceptual effect for changing the overall position (center-of-activity) of the pattern within sniff, possibly arising from weak modulation of glomerular activity by sniff-coupled mechanosensory responses (*14*). Alternatively, mice may be using the overall position of the pattern in sniff as a weak cue to solve the task, as the average randomly-generated Non-target pattern has a different position within sniff compared to the Target pattern.

We found a primacy effect where earlier-activated glomeruli have larger effects on perceptual responses. Primacy has been suggested as a strategy for animals to recognize the same odor across varying concentrations, as early-activated glomeruli remain stable across different concentrations. Animals trained to recognize odors across varying concentrations were impaired during coarse optogenetic disruption, but only when the disruption occurred in the initial ~100ms of inhalation (*15*). In the present study, animals were trained to recognize a single Target synthetic odor, and displayed a primacy effect without explicit training to ‘concentration-variants’ of the Target. Hence, the preferential weighting of early inputs occurs under different task demands, suggesting that primacy is a fundamental property of the olfactory system. Primacy may be supported by downstream computations, at the level of mitral cells (*35*) or piriform cortex (*16, 36*).

We cannot rule out the possibility that fibers of passage were activated during our spot stimulation. However, this effect is likely to be weak—MT cells were activated by few localized spots at the laser intensity chosen for our experiments. Another study also using OMP-ChR2 mice, with similar stimulation size, duration and intensity have also found negligible effects of fibers of passage (*11*). Increasing the stimulation intensity led to more apparent recruitment of fibers of passage, reflected in increased number of active spots located anteriorly (unpublished). Furthermore, we observed negligible non-linear interactions between spots during spatial replacements, which is not expected if there is widespread co-activation of glomeruli from fibres of passage.

Another limitation of the study is that our synthetic stimuli may not capture the full complexity of glomerular activity evoked by natural odors. We instead view the synthetic approach as complementary to approaches using more naturalistic stimuli. The claim is not that these synthetic stimuli are direct proxies for naturalistic stimuli—rather, synthetic stimuli afford well-controlled experiments with precise parameterization and causal manipulation, and can be used to establish basic principles of the neural code. In other well-studied sensory systems, foundational understanding of sensory processing has been built upon synthetic, reduced stimuli (*37*), while naturalistic stimuli have been used to test and refine foundational models (*38*). Future experiments may test the STM model by measuring perceptual distances between large sets of chemical odors while recording calcium responses to these odors in parallel.

Future studies may elucidate the specific circuit mechanisms for spatio-temporal template matching, by recording MT cells or piriform cortex responses to parametric patterned stimulation. Specific parameters or operations in the model may eventually be mapped to specific circuit computations. For example, the time constant in waveform activation, may strongly depend on rapid inhibition of MT cell activity after initial excitation (*13, 39*). The time constant for primacy may depend on a combination of recurrent inhibition in MT and cortical circuits (*16, 35*). Template matching procedures may be implemented by biologically-plausible circuits that implement delay lines (*40, 41*).

We developed a novel experimental and theoretical approach that links the complex spatio-temporal language of the brain with perception and behavior. This is especially pertinent, given continued advancements in optical techniques that allow direct access and manipulation of the spatiotemporal neural codes, at the level of computations consequential to behavior (*42–45*).

## Acknowledgments

We thank A. Resulaj, G. Serrano, A. Pickens, S. Stark, S. Toole, H. Ma, G. Lerman, J. Kappel, N. Amin, A. Oganov, M. Robles-Long, and A. Pillai for assistance in building and running experiments. We also thank B. Mensh, K. Nagel, T. Movshon, W. Ma, J. Gill and H. Nakayama for helpful comments.

This work was supported by the NIH BRAIN Initiative Grant R01NS109961.

## Author contribution

E.C., S.P., S.S. and D.R. conceived and designed the experimental, computational and modeling approaches; E.C. and C.W. built the experimental system and performed experiments; M.M. and E.C. performed computational analyses; E.C., M.M., S.S., S.P., and D.R. wrote the manuscript; S.P. and D.R. supervised the project.

## Methods

### Mice

Behavior and electrophysiology experiments were conducted in OMP-ChR2-YFP heterozygous mice. Control experiments were conducted in B6(Cg)-Tyrc2J/J (albino B6) mice (Jackson labs). Subjects were 8–12 weeks old at implantation and were maintained on 12hr light–dark cycle in isolated cages after implantation. All procedures were approved by the IACUC of NYULMC in compliance with the NIH guidelines for the care and use of laboratory animals.

### Surgery

Mice were anesthetized with isoflurane during surgical implantation (2.0% during induction, 1.5% during surgery). A circular craniotomy was performed to expose both hemispheres of the dorsal olfactory bulb (3 mm craniotomy extending from the rostral rhinal vein to the naso-frontal suture, centered on the midline) using either an air-driven dental drill (Midwest Tradition, FG 1/8 drill bit), or a 3mm biopsy punch (Miltex). A cranial window was implanted, replacing a circular piece of skull by a glass coverslip (3 mm diameter, Warner Instruments) that was secured in place using a mix of self-curing resin (Orthojet, Lang Dental) and cyanoacrylate glue (Krazy Glue). For electrophysiological recordings, the cranial window contained two ~1 mm diameter holes pre-drilled and filled with a silicone elastomer (Kwik-Sil, World Precision Instruments). A custom 3D-printed headpost was placed around the cranial window and affixed to the skull using C&B Metabond dental cement (Parkell). Each animal recovered for at least 10 days prior to experiments.

### Patterned photostimulation

To activate olfactory bulb glomeruli in specific temporal sequences we used a digital micromirror device (DMD) projector system (Mightex Polygon 400). We also custom built a similar system, using a 473 nm fiber-coupled diode-pumped solid state laser (CNI Laser MBL-N-473, 1.5W). The fiber output was passed through a piezo-driven homogenizer (Mightex) to remove laser speckle, and collimated and expanded (Thorlabs). The beam was then modulated by a DMD (ALP-4.2, Vialux, Germany) to produce light patterns with 10 μm spatial and 1 ms temporal resolution. These light patterns were projected off a 90/10 beam-splitter (Thorlabs) into a 4x objective (Olympus) which focused the patterns onto the surface of the olfactory bulb.

### Mitral cell responses to spot stimulation

NeuroNexus A64 Poly5 2 × 32 probes were used to record acutely from two awake animals. Units were detected using Spyking Circus (v0.3.0). Single 120μm square spots were stimulated at 15mW/mm^2^ on the olfactory bulb with ~10 repetitions per spot, randomly interleaved within a single session. To construct response heatmaps, responses are represented as the change in firing rate during the stimulus onset (80ms) compared with each unit’s average firing rate within the session. PSTHs were smoothed with a 3-sigma Gaussian kernel of width 30 ms.

### Behavioral training

All behavioral events (sniff, stimulus delivery, water delivery, and lick detection) were monitored and controlled by custom programs written in Python interfacing with a custom behavioral control system (Janelia Research Campus) based on an Arduino Mega 2560 microcontroller. Behavioral experiments began after at least 7 days of water restriction (1ml/day). Mice were housed on a reverse light/dark cycle, and training took place during the day. To acclimatize animals to head-fixation and the behavioral setup, animals were shaped by water given through a single lick tube until they received their entire 1 ml water ration during a session. In subsequent sessions, a second lick tube was introduced. To encourage exploratory behavior in subsequent training, animals were rewarded for alternating licks between left and right lick tubes. Two-lick shaping sessions persisted until animals successfully received entire 1 ml water ration in a session. Licking was detected using a capacitive touch sensor (Sparkfun) coupled to hypodermic tubing which triggered the release of water droplets by a pinch valve. Sniff was monitored using a pressure transducer coupled to a custom Teflon odor port (https://github.com/c-wilson/olfactometry/tree/d885d09d5d9544bc746e732705ecb4618f15dbd0/parts/odor_ports).

Animals performed a 2-alternative forced choice task, in which ‘left lick’ and ‘right lick’ were randomly assigned to Target and Non-target patterns for each animal. Trials were randomly interleaved in each session (~600 trials/day). Each trial consisted of a stimulus period, a grace (600 ms), response period (1 s), and an inter-trial interval with variable duration (5-6 s). On all trials, masking LEDs signaled the start of the stimulus period and persisted through the stimulus period and delay. A broadband tone (piezo buzzer) signaled the start of the response period. The masking LEDs were chosen such that the central wavelength matched that of the optogenetic stimulation (473 nm) and were positioned near the animals’ eyes to prevent task performance based on visual cues. The animal’s choice is recorded as the first lick during the response window. The trial ends as soon as the animal makes a choice during the response window, or after the entire window duration has elapsed. During initial training, some mice displayed side-biased licking, consistent with previous reports in the literature (*46*). We employed a de-biasing procedure which would increase the incidence of trials on the biased-against side(*47*), and was successful at removing this bias and not used after the training period.

Each animal was first trained to discriminate one Target from one Non-target pattern, and after exceeding criterion performance of 80%, animals were trained to discriminate the same assigned Target from randomly-initialized Non-target patterns. Because of the combinatorial possibilities, Non-target patterns were never repeated. During initial training, correct choices were rewarded 100% of the time but the animal was gradually shaped towards 70% reinforcement. In test sessions, probe trials (Target perturbations) were introduced. Probe trials were never rewarded regardless of the animals’ responses, substituting a portion of the unrewarded Target or Non-target trials. Probe trials formed 10% of total trials while Targets and Non-targets formed 45% of trials. Olfactory experiments using a similar trial structure (learned versus probe stimuli) have been previously reported(*15, 48*).

We presented several different types of probe trials. In spatial perturbations, one or more Target spots were replaced with randomly selected Non-Target spots, while the activation timing of all spots was fixed and equal to the Target pattern activation timing. On any spatial perturbation trial, there was an equal probability of having *n* spots replaced where *n* ranges from 1 to 5. For a given *n* spot replacement, the combination of spots replaced was randomly chosen.

In random temporal perturbations, the spatial identity of stimulated spots was the same as the Target pattern, but the activation onset of one or more spots was changed, with spot duration unchanged. For single spot shifts, one randomly-chosen spot with latency *t*_*i*_ is shifted randomly by *Δt*_*i*_ ∈ {−100, −90, …, 0,10, …, 100} ms. Positive (negative) shifts indicate shifting the spot later (earlier) within sniff. Here and later, all temporal shifts are constrained such that the new stimulation time *t*_*i*_ + *Δt*_*i*_ does not precede inhalation (or *t*_*i*_ + *Δt*_*i*_ ≥ 0). For earlier activated spots in the pattern, this leads to a range of shift that is asymmetric. This constraint is due to the fact that our stimulation patterns are always triggered after inhalation is detected, and it would be practically difficult to initiate a pattern at some consistent defined duration before inhalation.

For random shifts of multiple spots simultaneously, we included various shift procedures. Because it is practically impossible to sample every combination of spot shifted and shift amount, we constrained our shifts in different ways. In one shift paradigm, each spot *i* is shifted randomly by *Δt*_*i*_ ∈ {−80, −70, …, 0,10, …, 80} ms. In another shift paradigm, we performed random scrambling of the order of spots while keeping the timing structure within the pattern fixed.

Because these random shifts tend to induce large amounts of shift in the pattern, we also introduced an additional shift paradigm biased towards smaller shifts: any spot *i* is shifted randomly by *Δt*_*i*_ ∈ {−40, −20, 0,20,40} ms with Σ_*i*_|*Δt*_*i*_| ≤ 180 ms. Finally, because of differential effects of shifting spot latencies relative to each other versus shifting the entire pattern synchronously, we also introduced a shift paradigm where a “global” shift *Δt*_*glob*_ ∈ {30,60} ms was applied equally to all spots, in addition to single spot shifts of *Δt*_*i*_ ∈ {−80, −70, …, 0,10, …, 80} ms. To summarize the overall amount of temporal shift in the pattern regardless of which combination of spot shifts occurred across all paradigms, we defined a value of 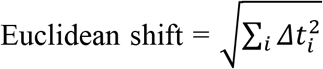.

In combined spatio-temporal perturbations, one, two, or three spots were first chosen to be replaced. The probability of replacements for one spot = 0.6, two spots = 0.3, three spots = 0.1. A separate set of spots were temporally shifted randomly by *Δt*_*i*_ ∈ {−80, −60, …, 20, …, 80} ms with *Δt*_*i*_ > 0. The probability of temporal shifts for one spot = 0.6, two spots = 0.3, three spots = 0.1.

### Detection thresholds for single spot stimulation

OMP-ChR2 mice and 2 control albino B6 mice were tested on a Go / NoGo task. Blue masking LEDs signaled the start of the trial and persisted through the stimulus period and response window (1s), with an inter-trial interval of 4-4.2 s. NoGo trials consisted of a randomly chosen single square spot (120μm, 80ms duration) over the olfactory bulb activated between 10 and 80ms after inhalation onset. If animals licked during the response period of a NoGo trial, the interval preceding the next trial was doubled to 8-8.4s as punishment. The intensity of stimulation was randomized across NoGo trials, chosen in steps between 2mW/mm^2^ and 50mW/mm^2^. Successful withholding of responses during NoGo trials was not rewarded. On Go trials, no optogenetic stimulation occurred, and animals were given water reward for a lick response. If animals did not lick, there was no further punishment.

### Perceptual distances for lowered laser intensity

OMP-ChR2 mice were trained to discriminate Target from Non-target patterns. For two mice, Targets comprised of two spots stimulated at 10 and 50 ms from inhalation, with Non-Targets containing two randomly-chosen, off-Target spots. For the other two mice, Targets comprised of four spots stimulated at 10, 50, 90 and 130 ms from inhalation, with Non-Targets containing four randomly-chosen, off-Target spots. The stimulation intensity used during training was 100mW/mm^2^, and all other stimulation parameters, shaping and training procedures, were identical to the main perceptual distance experiments. After training, Probe trials were introduced, which are Target patterns with different stimulation intensities, randomly chosen in integer increments from 0 to 255, with 255 corresponding to the intensity used during training and 0 corresponding to no stimulation.

### Logistic regression models

#### Quantifying spatial and temporal perturbations with logistic regression

To quantify the effects of spatial and temporal perturbations (Fig. 2) we fit logistic regression models (*49*) with linear terms. For spatial perturbations, we fit:

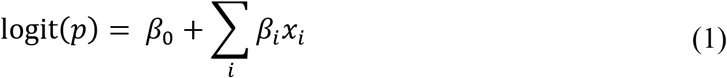

where *p* is the probability of making a like-Target response. *x*_*i*_ is a binary variable taking the value 1 during trials where spot *i* is replaced and 0 otherwise. Here and later, *β*_*i*_ is the weight on spot *i*, and *β*_0_ is the bias. For temporal perturbations, we fit:

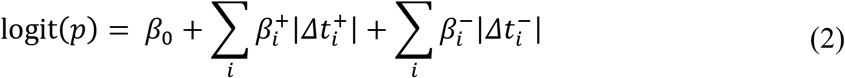

Where 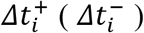 is the amount in ms that spot *i* has been shifted later (earlier) within the sniff cycle.

#### Regression models with interaction terms

To test the prediction that a linear readout of single spot effects is sufficient to account for spatial perturbations, we tested the linear model (Equation 1) against a model that allows for non-linear interactions between spots during replacement (Fig. 5C,D):

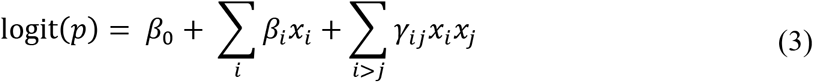

The model is similar to Equation 1, with an additional weighted term *x*_*i*_*x*_*j*_, which equals 1 if both spots *i* and *j* were replaced on the same trial, and 0 otherwise.

To test the prediction of non-linear temporal interactions between spots, we fitted a logistic regression model that allows for non-linear interactions between spots during temporal shifts (Fig. 5E), which we refer to as *Δt* + paired shifts model:

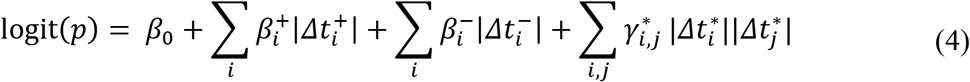

The model is similar to Equation 2, with an additional weighted term 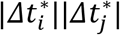, which is the magnitude of temporal shift in spot *i* multipled by the magnitude of temporal shift in spot *j*, in each unique combination of positive or negative shift in spot *i* and *j*, with corresponding coefficient 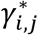. A positive (negative) coefficient implies that the effect of shifting both spots in their specified directions is larger (smaller) than predicted from the sum of effects from shifting either spot individually—super-linear (sub-linear) additivity.

#### Alternative temporal coding regression models

Center-of-latencies (*T*_*L*_ model):

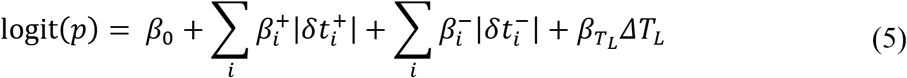

With the center-of-latencies of the pattern defined as 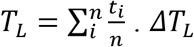 is the difference in center-of-latencies of the probe pattern compared to the Target pattern. 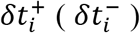 is the amount in ms that spot *i* has been shifted later (earlier) within the sniff cycle, but for latency defined relative to *T*_*L*_. For example, if spot *i* in the Target pattern occurred 20 ms before 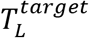, and the same spot in a probe pattern occurred 40 ms later than 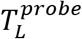, then 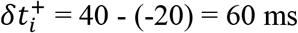.

*Δt* + paired time:

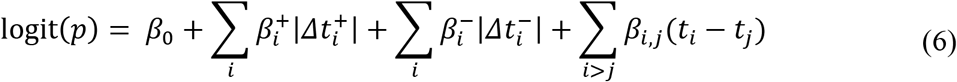

Similar to Equation (2) but with additional *t*_*i*_ − *t*_*i*_ that considers the relative latencies between all pairs of spots.

*Δt* + reduced pairs:

Same as Equation (6), but only consider *t*_*i*_ − *t*_*i*_ where (*i, j*) ∈ {(1,2), (2,3), (3,4), (4,5), (5,6)}.

*T*_*L*_ model + reduced pairs:

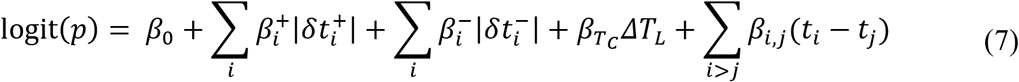

Same as Equation (5) but with additional *t*_*i*_ − *t*_*j*_, for (*i, j*) ∈ {(1,2), (2,3), (3,4), (4,5), (5,6)}.

Rank order:

We consider a model of the rank order of spots, ignoring the exact latencies of activation. We denote with *R*_*i*_ the ranking of spot *i* in the Target pattern and with *Z*_*i*_ the ranking of the same spot in the considered probe pattern. In case of ties, we average *Z*_*i*_ across simultaneously activated spots. The predictor relative to each spot is the change in its rank order, computed as *dr*_*i*_ = (*R*_*i*_ − *Z*_*i*_)^2^

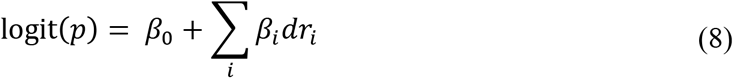

Null model:

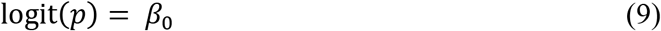

### Spatio-temporal Template Matching (STM) model

We define neural activation space as a *N*-dimensional space, where *N* is the number of channels (neurons, or glomeruli, or activated spots) conveying neural signals. We represent a spatio-temporal pattern of neural activity *P* as a (*N* ∗ *2*)-dimensional array:

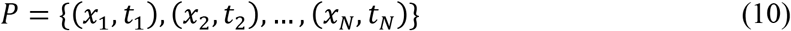

Where *x*_*i*_ is a binary variable indicating whether the *i*-th channel is active, and *t*_*i*_ is the activation onset of the *i*-th channel. If the *i*-th channel is not active, then we set *t*_*i*_ to 0.

Given *P* that is defined as a discrete object, we map it onto a set of continuous waveforms, applying the convolution with an exponential decaying kernel *h*(*t*; τ_*act*_) defined as:

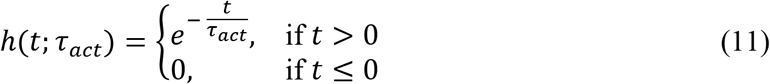

with τ_*act*_ > 0.

The mapping to continuous functions through convolution is defined as:

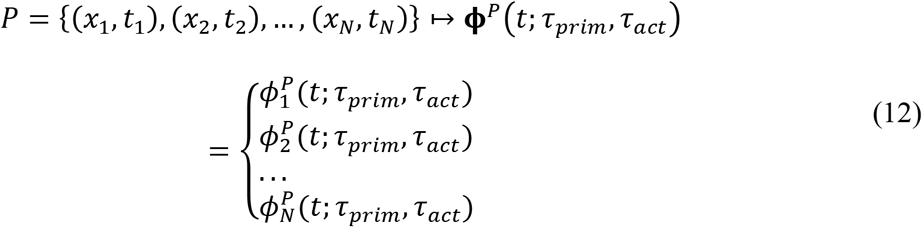

where

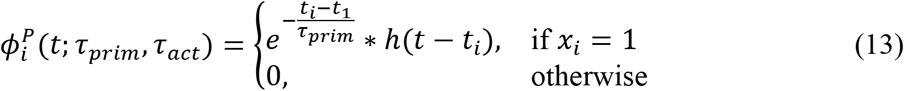

and τ_*prim*_ > 0.

Given an angle 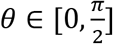, we consider in the N-dimensional space N axes rotated by an angle *θ*. We denote with **e**_*i*_(*θ*) the direction of the *i*-th axis and project the components of the function **ϕ**^**P**^ on these axes:

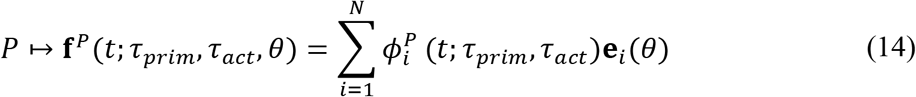

The angle *θ* can be interpreted as a parameter that represents the relevance of channel identity in the readout. For example if, *θ* = 0, then channel identity is not relevant and signals from all channels are summed up before within-channel comparisons. If 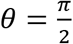, then each channel is considered independently. For our model fitting, we set 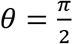. We have alternatively fit *θ* free parameter, finding *θ* close to 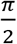.

Once the patterns are represented through continuous functions, we assign to each pattern *P* its center-of-activity. We defined the center-of-activity *T*_*C*_ as the timepoint which covers half of the waveform area in *P*:

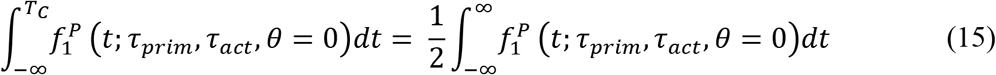

The center-of-activity essentially represents the overall position of the pattern within the sniff cycle. Alternative definitions of center-of-activity yield comparable or poorer fits, such as by considering the mean (center-of-mass) of the waveform area, the center-of-latencies or just the latency of the earliest-activated spot in the pattern.

Given a target pattern *P* and a probe pattern *Q*, we define the perceptual distance between *P* and *Q* in the following steps:

1. Calculate for each pattern its center-of-activity, *T*_*C*_
2. Compute the difference between the centers of activity of the two patterns: 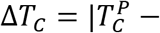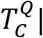.
3. Align the patterns to their center-of-activity. We denote with *P*′ the shifted pattern

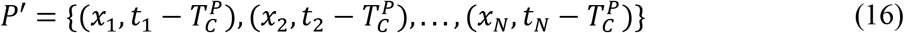

And we do the same for *Q*′.
4. The perceptual distance between the two patterns is given by:

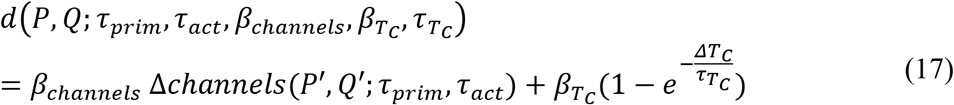

Where

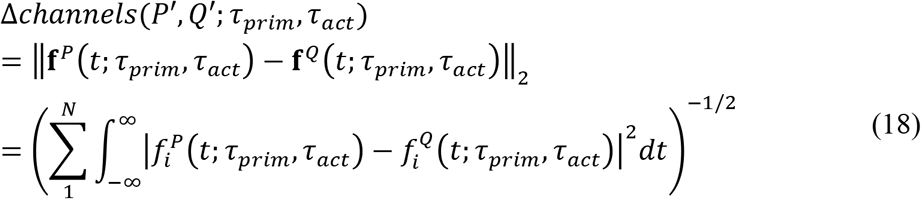

with 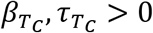.

The animal’s probability *p* of making a like-Target response to any given pattern then decreases with increasing perceptual distance *d*, given by the logistic function on *d*:

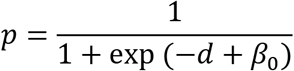

Where *β*_0_ is the bias term.

### Comparison of STM model with regression

#### Temporal interactions

For quantifying temporal interactions produced by the STM model, we first simulated a temporal shift dataset of 400,000 trials starting from Target patterns with the same timing structure as in the behavioral experiment. On each simulated trial, each spot *i* is shifted randomly by *Δt*_*i*_ ∈ {−100, −99, …, 0,1, … 99,100} ms such that *t*_*i*_ + *Δt*_*i*_ ≥ 0.

The responses of the STM model to temporally-perturbed patterns are then fitted with a logistic regression model allowing for temporal interactions (Equation 4). The regression coefficients are then compared to the same regression coefficients obtained by fitting to animals’ responses for all temporal perturbation data excluding synchronous shifts (Fig. 5E). Both sets of correlation coefficients were also compared by with Pearson’s correlation.

#### STM model versus regressions on all data

We compared the predictions of the STM model versus regression. For predictions on Non-target patterns and all Target perturbations except synchronous temporal shifts, the fitted regression model was a combination of the linear spatial and temporal models (Equation 1 and 4):

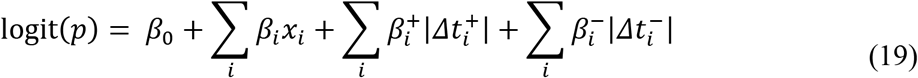

For predictions on synchronous temporal shifts, the fitted regression model was

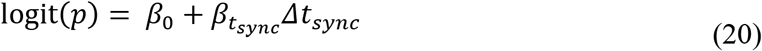

Where *Δt*_*sync*_ is the amount of synchronous shift.

### Model fitting and evaluation

#### Logistic regression models

To quantify the effect of individual spatial or temporal perturbations (Fig. 2), we fit regression models using statsmodels in Python (*50*). We also implemented the same fits using glmnet in R (*51*), with elastic net regression and the elastic net parameter set to 0.5, yielding similar results. All other fits were performed with glmnet. For any model comparison, a cross-validation procedure was performed. To compare between spatial regression models, 5-fold cross-validation was performed on all spatial perturbation trials plus Target and Non-target trials, from the dataset from all animals pooled together. For comparing regression models to the STM model, the full dataset was split into a training set (75% of total trials) and a test set (25% of total trials). To fit each model, we performed 5-fold cross-validation on the training set. The proportion of each trial type (Target, Non-Target and each type of Probe trial) was kept the same within each fold and the test set to match the proportion in the overall dataset. The same splits were used across all models we considered. To compare among temporal regression models (Fig. S2), the same splitting procedure was used, but restricted to temporal perturbations only. We selected as best-fitting parameters those parameters that returned the lowest cross-validated prediction error (measured as percentage of incorrectly predicted trials) averaged across folds. We then evaluated the accuracy of each model by computing the Brier Score (BS) on the test set.

For a binary variable *x*, the Brier Score is defined as

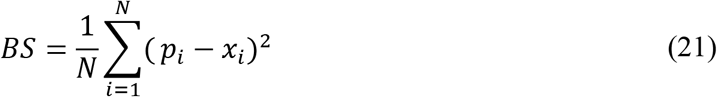

where *N* is the number of samples, *p*_*i*_ is the predicted probability of a positive outcome in the *i*-th sample and *x*_*i*_ is the observed outcome in the *i*-th sample. BS ranges between 0 (perfect prediction of the variable outcome) and 1 (totally wrong prediction). BS is a commonly accepted standard for evaluating predictions of binary outcomes, and preferred over measuring the fraction of correctly classified responses (*52*). In our case, alternatively measuring the percentage of correct binary classifications did not change our conclusions.

#### STM model

To fit the STM model we used the same training and test sets and splitting folds as for logistic regression. We defined a grid in the 5-dimensional parameter space and, for each grid point, we computed perceptual distances on the training set trials. Perceptual distances are a function of Δ*channels*, Δ*T*_*C*_ and the bias and the bias term. We used logistic regression to find their relative weights *β*_*channels*_, 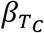 and *β*_*0*_ in predicting behavioral choice under a 5-fold cross-validation procedure. The parameters for the logistic regression were obtained using the glmnet package in R (elastic net penalty parameter = 0.5). We considered as best fitting parameters for the perceptual distance those parameters in the grid that returned the lowest prediction error averaged across folds. For these parameters we ran again logistic regression on the total training set to estimate the final logistic regression weights. We evaluated the goodness of fit of the perceptual metric model and the comparison across different models using the test set in the same way we did for logistic regression. This grid-search procedure is a brute-force approach and does not guarantee the convergence to the optimal solution. Despite this, the fitted STM model performs as well as the separate logistic regression fits which are optimized solutions.

#### Model comparison

To perform statistical tests on model comparisons, we first computed the Brier Score on a *N* = 500 bootstrapped version of the test set. For pairwise model comparisons, we performed a t-test on the Brier scores. For comparing between variants of the STM model (Fig. 5B) or variants of temporal coding models (Fig. S2), we compared models using ANOVA and post hoc tests corrected for multiple comparisons (Tukey's Honestly Significant Difference Procedure). All statistical tests were performed in MATLAB.

## Supplemental Figures

**Figure S1.**
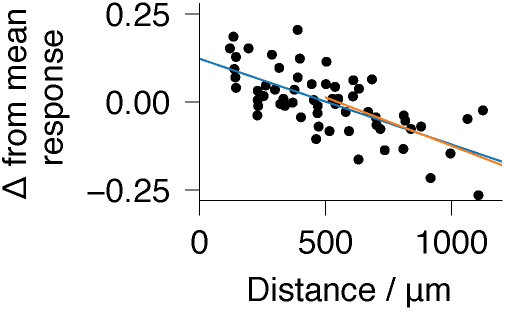
Effect of spot distances on spatial replacements. Single spot replacement trials were grouped by spot position (1 through 6) replaced. Within each group, trials were further subdivided (ten bins) by the spatial distance between the position of the Target spot being replaced, and the random replacing spot. Responses per bin were then normalized to the mean across all bins (i.e. mean response by spot position), and plotted as a single point. Linear regression was used to fit all the data (blue, R^2^ = 0.46, p < .001), or separately to trials with distance > 500μm (orange, R^2^ = 0.35, p < .001). Because the perceptual effect of distance was true both at small and large distances, the effect cannot be accounted for by the alternative explanation that adjacent spots directly stimulate shared glomeruli.

**Figure S2.**
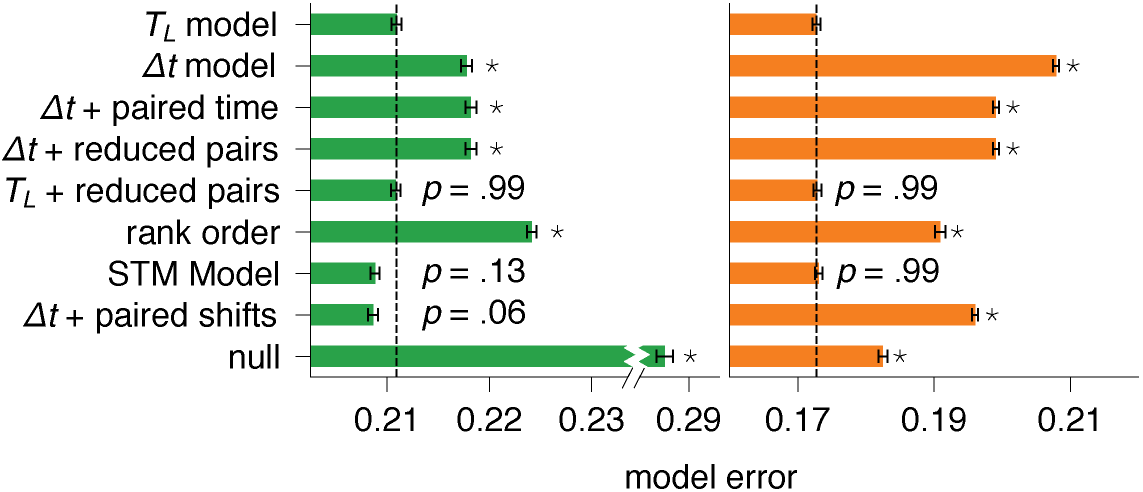
Comparisons across different models of timing (Methods): a centroid model that encodes center-of-latencies *T*_*L*_ and individual spot latencies relative to *T*_*L*_, is compared with alternative models in predicting responses to temporal perturbations, on all temporal shift trials excluding synchronous shifts (green) and only for synchronous shifts (orange). Dashed line indicates mean error for *T*_*L*_ model. Asterisks indicate p < .001 significant difference from *T*_*L*_ model, corrected for multiple comparisons. The STM model, which computes centroids over template waveforms, is also shown.

**Figure S3.**
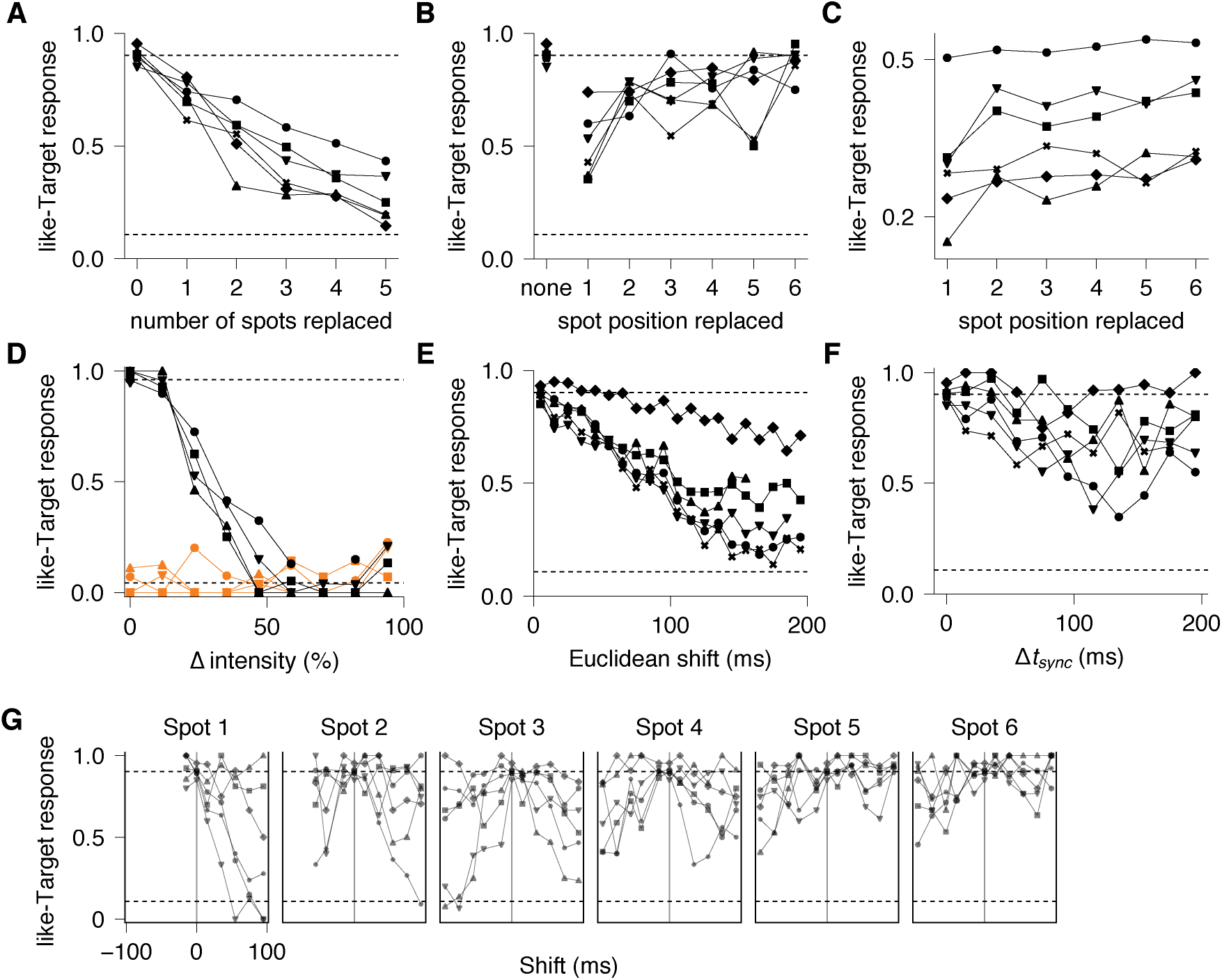
Individual mouse responses to Target perturbations. **(A)** Increasing the number of spots replaced. **(B)** Changing spot position replaced in single spot replacement. **(C)** Changing spot position in multiple spot replacement. **(D)** Changing stimulation intensity. **(E)** Increasing Euclidean shift. **(F)** Increasing synchronous shift. **(G)** Single spot shifts.

